# When two cells are better than one: specialized stellate cells provide a privileged route for uniquely rapid water flux in *Drosophila* renal tubule

**DOI:** 10.1101/763664

**Authors:** Pablo Cabrero, Selim Terhzaz, Anthony J. Dornan, Saurav Ghimire, Heather L. Holmes, Daniel R. Turin, Michael F. Romero, Shireen A. Davies, Julian A. T. Dow

**Affiliations:** Institute of Molecular, Cell and Systems Biology, College of Medical, Veterinary and Life Sciences, University of Glasgow, Glasgow G12 8QQ, UK; Department of Physiology and Biomedical Engineering, Mayo Clinic College of Medicine & Science, Rochester, MN 55905; Division of Nephrology and Hypertension, Mayo Clinic College of Medicine & Science, Rochester, MN 55905; University of Minnesota Rochester, Rochester, MN 55905

**Keywords:** Malpighian tubule, *Drosophila melanogaster*, aquaporin, *Xenopus* oocyte, stellate cell

## Abstract

Insects are highly successful, in part through an excellent ability to osmoregulate. The renal (Malpighian) tubules can secrete fluid faster on a per-cell basis than any other epithelium, but the route for these remarkable water fluxes has not been established. In *Drosophila melanogaster*, we show that 4 members of the Major Intrinsic Protein family are expressed at very high level in the fly renal tissue; the aquaporins Drip and Prip, and the aquaglyceroporins Eglp2 and Eglp4. As predicted from their structure and by their transport function by expressing these proteins in *Xenopus* oocytes, Drip, Prip and Eglp2 show significant and specific water permeability, whereas Eglp2 and Eglp4 show very high permeability to glycerol and urea. Knockdowns of any of these genes impacts tubule performance resulting in impaired hormone-induced fluid secretion. The *Drosophila* tubule has two main secretory cell types: active cation-transporting principal cells with the aquaglyceroporins localize to opposite plasma membranes and small stellate cells, the site of the chloride shunt conductance, with these aquaporins localising to opposite plasma membranes. This suggests a model in which cations are pumped by the principal cells, causing chloride to follow through the stellate cells in order to balance the charge. As a consequence, osmotically obliged water follows through the stellate cells. Consistent with this model, fluorescently labelled dextran, an *in vivo* marker of membrane water permeability, is trapped in the basal infoldings of the stellate cells after kinin diuretic peptide stimulation, confirming that these cells provide the major route for transepithelial water flux. The spatial segregation of these components of epithelial water transport may help to explain the unique success of the higher insects.

**Significance statement:** The tiny insect renal (Malpighian) tubule can transport fluid at unparalleled speed, suggesting unique specialisations. Here we show that strategic allocation of Major Intrinsic Proteins (MIPs) to specific cells within the polarized tubule allow the separation of metabolically intense active cation transport from chloride and water conductance. This body plan is general to at least many higher insects, providing a clue to the unique success of the class Insecta.

## Introduction

There are more species of insects than all other forms of life combined. In part, this is because of the exceptional ability of the simple body plan to operate in a wide range of environments, and osmoregulation is a key component of this success. Remarkably, the insect Malpighian (renal) tubule is capable of secreting fluid faster (on a per cell volume basis) than any other epithelium known (1, 2). In *Drosophila*, the renal tubule has two major cell types (3-5); the mitochondria-rich principal cell actively transports protons via an apical, plasma membrane V-ATPase (6), setting up a gradient which is exchanged primarily for potassium (7, 8) and enters the cell basolaterally through a combination of Na^+^, K^+^-ATPase (9), potassium channels (10, 11) and cotransports (12-14). The smaller stellate cell (15, 16) provides a route for hormone-stimulated (17-20) chloride conductance through a basolateral ClC-a chloride channel (21), and secCl, an apical cys-loop chloride channel (22), to balance the lumen-positive charge, and so effect a net movement of salt. Aquaporins (e.g., the water transporting MIP proteins) are known to be highly expressed in insect tubules (23-27), and global knockdown of an aquaporin in the *Aedes* mosquito (28-30), or in the beetle *Tribolium* (31) impacts water loss. However, the route or mechanism of the very high osmotically-obliged water fluxes that produce such remarkable fluid output has not been characterised. Here, using the powerful cell-specific transgenic technologies unique to *Drosophila melanogaster* (32), we show that this flux is transcellular, and selectively through the stellate cells, mediated by two aquaporins, in response to diuretic hormone stimulation.

## Results & Discussion

### Tubules express 4 members of the Major Intrinsic Protein family

Major intrinsic proteins (MIPs) are a multigene family of 6-transmembrane domain proteins, that assemble as tetramers to form pores (33). Most members of the family are true water channels (aquaporins); others can facilitate movement of water or small organic molecules (aquaglyceroporins); and the substrates of some are still obscure (33). In *Drosophila*, eight genes make up the MIP family **(Fig. 1)**, but the FlyAtlas and FlyAtlas2 gene expression online resources (25, 34, 35) independently report that only four are expressed at high levels in epithelia such as the salivary gland, midgut, hindgut and Malpighian tubules **(Fig. 1).** Two of these highly expressed genes (*Drip and Prip*) are similar to classical aquaporins in structure, whereas the other two (*Eglp2* and *Eglp4*) align with the aquaglyceroporins (36). Comparison of the protein sequence of *Drosophila melanogaster* aquaporins (Drip and Prip) and aquaglyceroprins (Eglp2 and Eglp4) in a Clustal Omega alignment shows that key active-site residues, including those required for water selectivity and those involved for their regulation, have been conserved **(Fig. S1).** There are thus at least two candidates that could mediate high water flux rates in polarized epithelia.

**Fig. 1.**
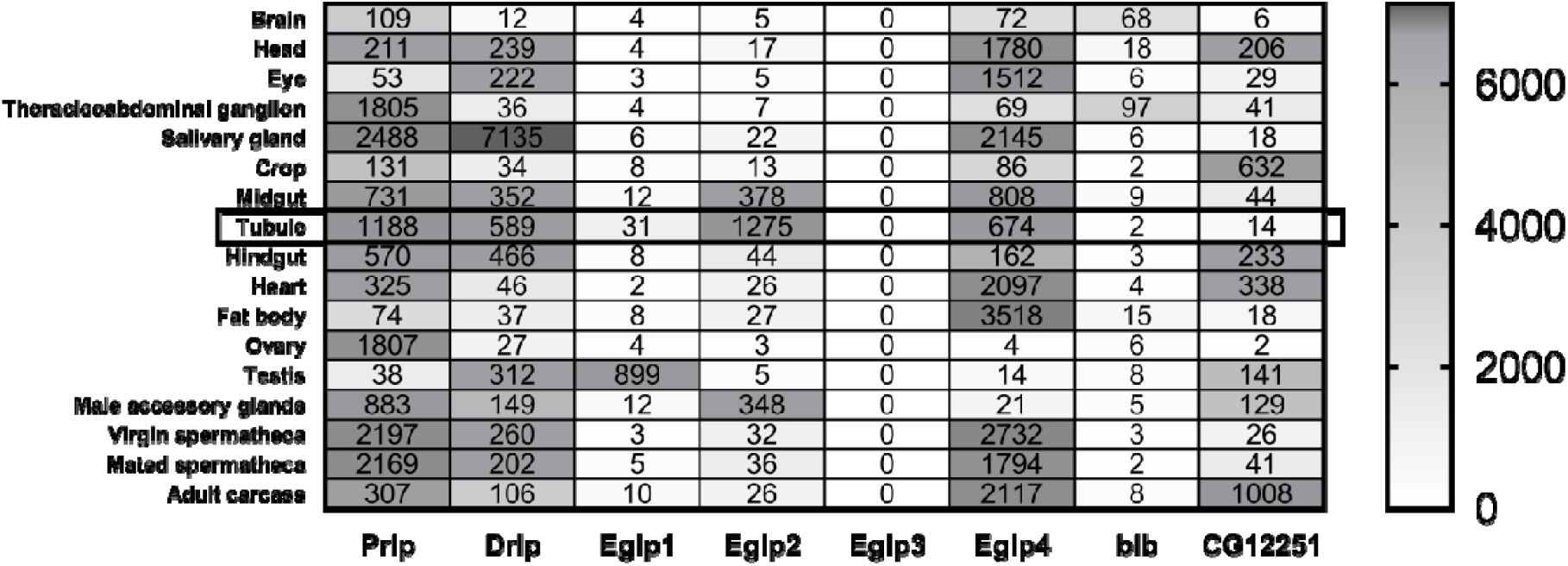
Major Intrinsic Protein (MIP) family expression in *Drosophila melanogaster*. Data mining of FlyAtlas.org identified four MIP genes *(Prip, Drip, Eglp2* and *Eglp4)* with highly abundant expression in adult Malpighian tubules.

### Each MIP localises to a different membrane domain within the tubule

Water and solutes transport are achieved by an apicobasally polarized distribution of membrane proteins, and accordingly, it is important to establish where in the tubule principal and stellate cells MIPs reside. We raised specific antibody against the four tubule-expressed MIPs and validated them by Western blotting (**Fig. S2 and Fig. S3**). Immunocytochemistry showed clear segregation of MIPs expression, with the two aquaporins expressed on opposite sides of the specialized stellate cell (Drip and Prip are localised to the basolateral and apical membranes respectively, **Fig. 2 *A* and *B*)**, and the two aquaglyceroporins on opposite sides of the main principal cell (Eglp2 and Eglp4 are localised to the basolateral and apical membranes respectively, **Fig. 2 *C* and *D***). Accordingly, overexpression of all 4 MIPs label with Venus (eYFP) recapitulate the pattern of expression observed by immunocytochemistry (**Fig. 2 *A’-D’***). These data are consistent with other reports that Drip and Prip show spatial separation in other insects, such as silkworm (37). It would thus be tempting to surmise that the stellate cell provides a major route for water flux through the tissue; but only the transport properties of one of these MIPs (Drip) has been established (23); how many of them are in fact functional aquaporins?

**Fig. 2.**
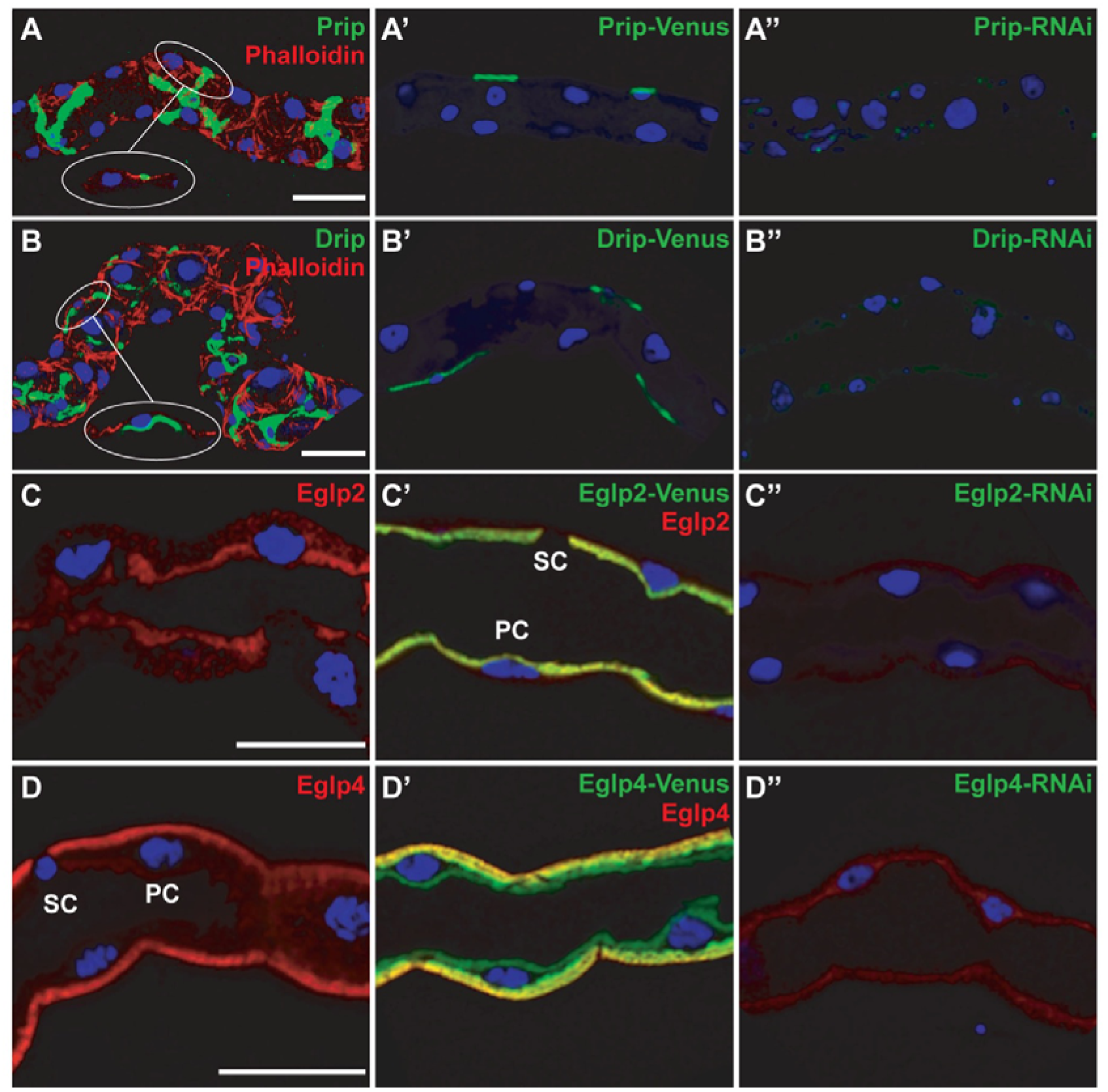
Subcellular localisation of MIPs in the *Drosophila* tubule. (*A, B)* Prip and Drip localise to opposite plasma membranes of the stellate cell (SC). (*A*) Prip is expressed in basolateral membrane (green) and is in a contact with the outside of the tubule, or hemolymph and (*B*) Drip in apical membrane (green) and faces the lumen of the tubule. DAPI (blue) staining for nuclei and phalloidin (red) staining for actin are shown. Inserts are single focal plane images showing magnifications of selected regions. (*A’, B’*) Overexpression of Prip and Drip label with Venus (eYFP) recapitulate the pattern of expression observed by immunocytochemistry. (*C, D*) Eglp2 and Eglp4 localise to opposite plasma membranes of the principal cell (PC). (*C*) Eglp2 is expressed in apical membrane (red) and (D) Eglp4 in basolateral membrane (red), and DAPI (blue). (*C’, D’*) Colocalization (yellow) between (*C’*) Eglp2-Venus and Eglp2 to the apical membrane and (*D’*) between Eglp4-Venus and Eglp4 to the basolateral membrane. (*A’’-D’’*) Down-regulation of MIPs in specific cell types using RNAi reduce protein levels. Scale bar = 40 µm.

### Stellate cell MIPs are aquaporins; Principal cell MIPs are aquaglyceroporins

Each of the four candidate genes was expressed in *Xenopus* oocytes, and tested both for classical swelling under hypoosmotic stress, and for facilitated flux of organic solutes. The two channels expressed in tubules (Drip and Prip) both acted as classical aquaporins, showing rapid water fluxes, but only barely detectable fluxes of organic solutes (**Fig. 3 *A* and *B*)**. By contrast, the Eglp2 and Eglp4 channels showed more modest fluxes of water, but rapid fluxes of small organic solutes, such as glycerol and urea, consistent with their predicted classification as aquaglyceroporins {Drake, 2015 #8090}. These data are thus in agreement with Drip and Prip providing a transcellular route for water through the stellate cells, and as the tubule provides a range of physiological readouts, this prediction can be tested experimentally.

**Fig. 3.**
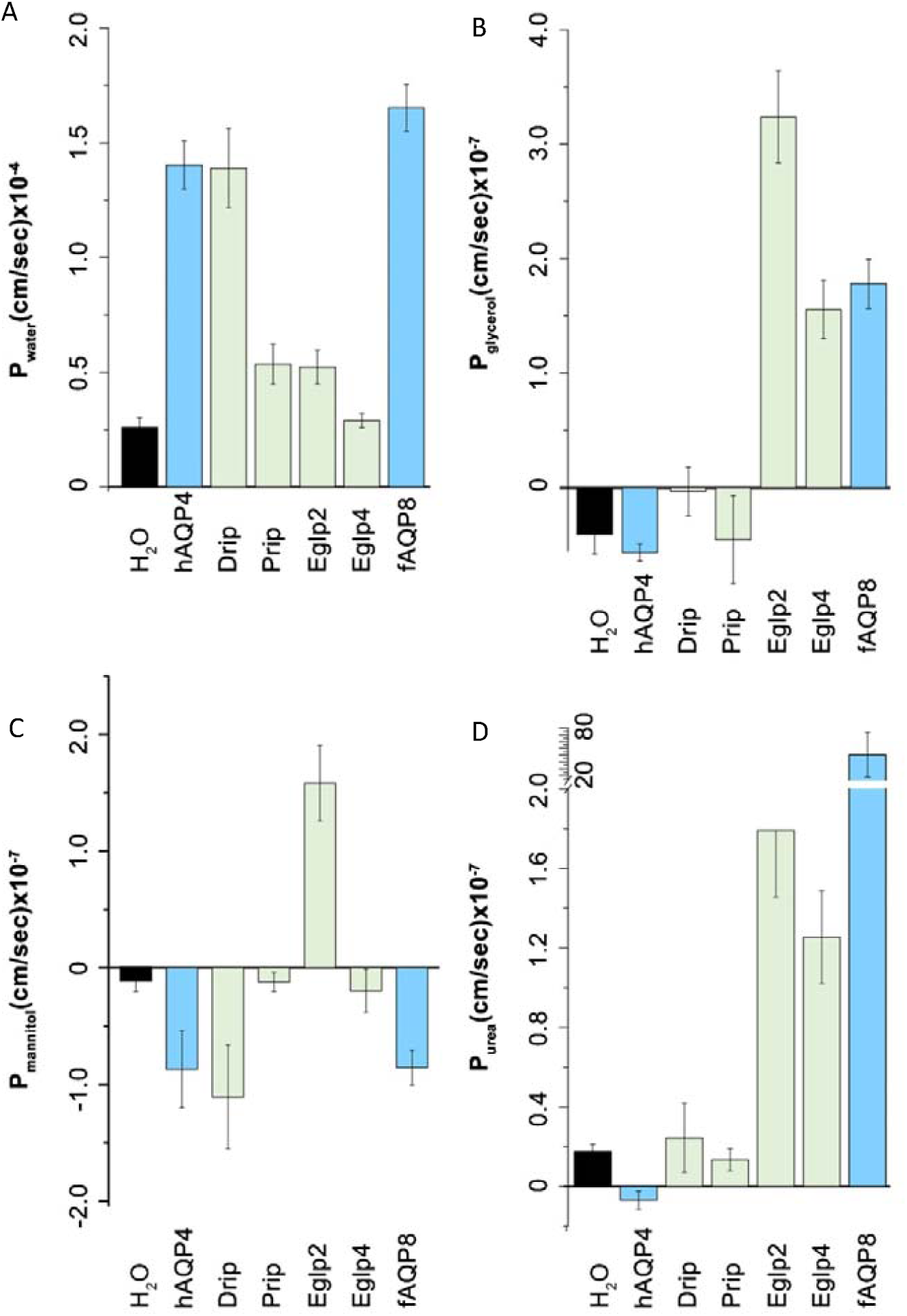
Transport specificity of *Drosophila t*ubule-enriched MIPs. Water-injected control oocytes or oocytes expressing *Drosophila* MIPs (Drip, Prip, Eglp2, Eglp4), human AQP4 (hAQP4) and mefugu AQP8 (fAQP8) were tested for permeability of (A) low osmolarity (Pf, P_H2O_), (B) urea (P_urea_), (C) glycerol (P_glycerol_) or (D) mannitol (P_mannitol_).

### Knockdown of AQPs reduces fluid transport and impacts survival

Although epithelial polarisation of some AQPs has been shown in other insects (23, 26, 37, 38), *Drosophila* genetic technology allows their physiological roles to be dissected with great precision. Using the GAL4/UAS system and renal cell-type specific drivers, it is possible to generate transgenic flies in which a single candidate gene is knocked down in only the tubule cell type in which it is expressed, leaving expression throughout the rest of the fly untouched. Accordingly, each of the four genes was knocked down in the cell-type in which their proteins had been shown to be expressed, and we were able to confirm by qPCR and immunocytochemistry the efficiency of the knockdown of MIPs expression at the gene and protein levels **(Fig. 4 *A* and 2 *A’-D’***). The resulting fluid output was then measured under baseline conditions, and when maximally stimulated with diuretic peptides of the capa and kinin families. Knockdown of either *Drip or Prip* in just the stellate cell significantly impeded fluid secretion, confirming functional roles in rapid fluid movement across the tissue (**Fig. 4 *B*).** However, knockdown of *Eglp2* or *Eglp4* in the principal cells also elicited reduced fluid secretion rates (**Fig. 4 *B***). This suggested two possibilities; either that all four MIPs could produce water conductance, through both cell types (at variance with the biophysical characterisation, **Fig. 3**), or that one pair of channels provided the main route for water, while the other pair allowed flux of an organic osmolyte, or metabolic substrate, such that blockade could reduce overall function of the tissue.

**Fig. 4.**
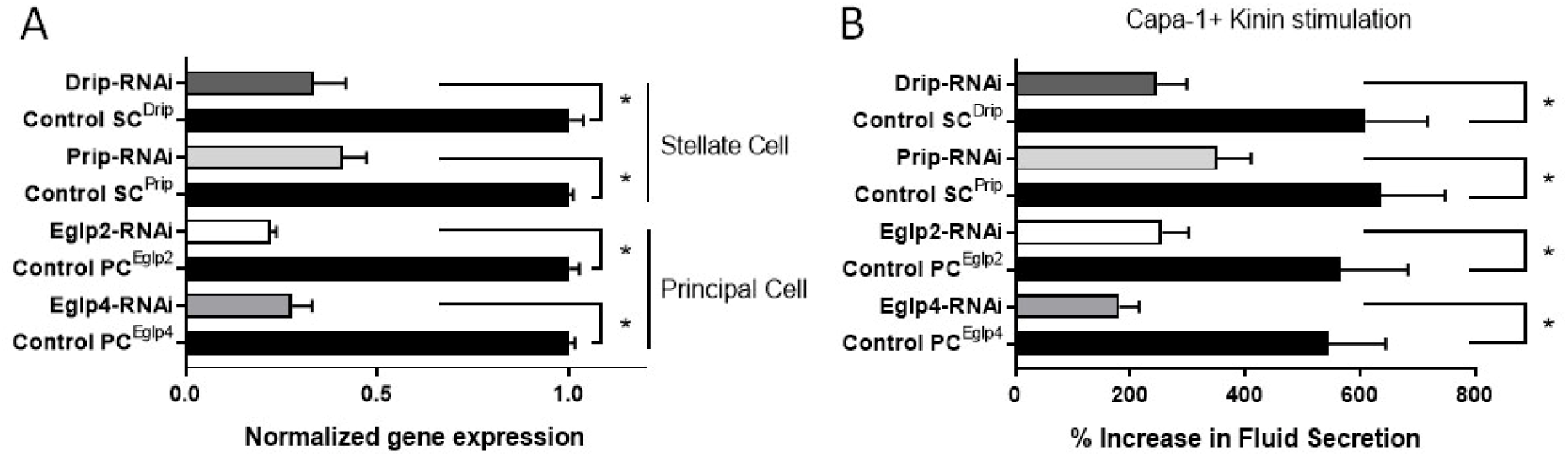
Validation of MIPs knockdowns and impact of cell-specific down-regulation of MIPs on fluid secretion. (*A*) Effects of knockdowns on tubule mRNA levels for MIPs, validated by qPCR. Cell-specific down-regulation of Eglp2 and Eglp4 in principal cells and Drip and Prip in stellate cells using their respective UAS-dsRNA lines. Data are expressed as mean fold change compared to parental controls ± SEM (*n* = 3). *P < 0.05 (Student’s t-test). (*B*) Impact of cell-specific knockdowns of MIPs on stimulated fluid secretion by tubules in response to Capa-1 and Kinin at 10^−7^ M. Data are expressed as percentage increase from basal fluid secretion compared to parental controls ± SEM (*n* = 6-10). *P < 0.05 (Student’s *t*-test).

The very high rates of generation of primary urine by the tubule could become a liability under dry conditions, and so knockdown of aquaporins would be predicted to impact survival under desiccation. This was shown by global knockdown of the *Drip* orthologue in *Anopheles gambiae*, the malaria vector (26); however, *Drip* is broadly expressed, and so the effect could not be attributed to the tubules (28, 29). Using GAL4/UAS technology, we were able to knock down *Drip* or Prip expression in just the tubule stellate cells, and show that this was sufficient to produce enhanced survival under desiccation stress **(Fig. S4).** Water flux across the tubule is thus limiting for terrestrial insects under desiccation stress, as previously suggested (39).

### The route of water flux is through the stellate cells

To distinguish the roles of the aquaglyceroporins from the aquaporins, it would be necessary to determine the route of water flux through the tubule. The complex polyglucan, dextran, can be readily fluorescently labelled, and also can be size-selected to ranges that can be swept along by water flux, but then trapped in a pathway of restricted permeability. Both the principal and stellate cells have apical microvilli, which in principal cells are stabilised by Fas2 (40) and contain mitochondria to support intense activity of the V-ATPase (41); and both cell types also possess basal infoldings, that increase the available surface area for transport (42). We thus stimulated tubules in the presence of fluorescently-labelled dextran, which pilot experiments had shown was too large to move across the epithelium. Dextran would thus accumulate in a compartment diagnostic of the route of water movement, be it the principal or stellate cells, or the paracellular route between the tight (‘septate’) junctions (43). The results showed that only the basal labyrinths of the stellate cells became labelled with 40 kDa dextran (**Fig. 5 *A* and *C*)**, and that the percentage of stellate cell population displaying uptake of dextran was significantly higher after kinin stimulation **(Fig. 5 *C***), thereby implying that the pathway provided by Prip and Drip in the stellate cells is the major route for water movement through the tubule.

**Fig. 5.**
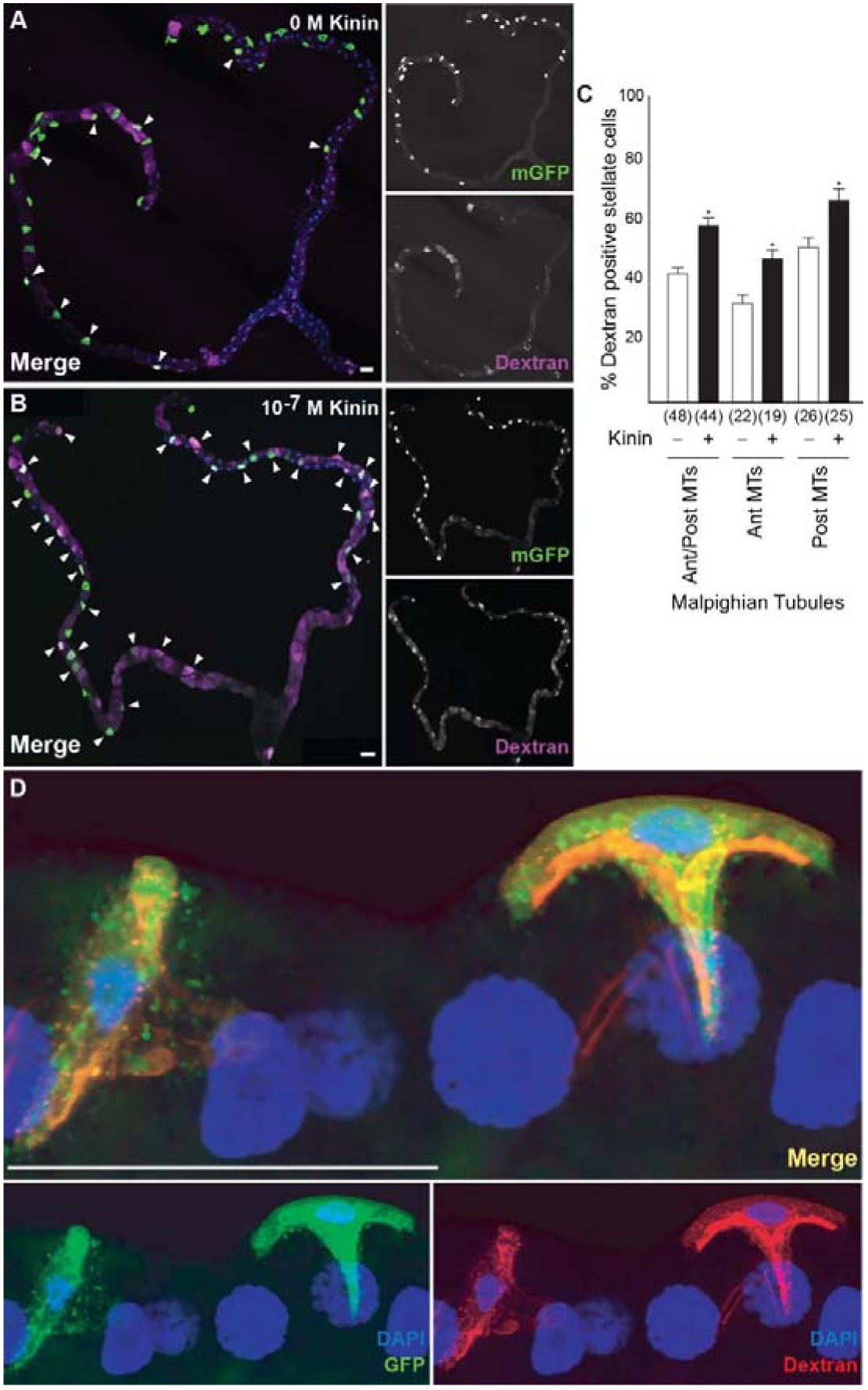
Dextran labelling demonstrates water flux specifically to the stellate cells. Accumulation of dextran of (*A*) unstimulated and (*B*) following application of Kinin (10^−7^ M) in tubule expressing mGFP in stellate cells. (*C*) Quantification of dextran labelling. Data are expressed as percentage of dextran positive stellate cells in response to 10^−7^ M Kinin compared to unstimulated tubule ± SEM (*n* = 44-48). *P < 0.05 (Student’s t-test). (*D*) Application of 40 kDa dextran conjugated to TRITC dye (red) to tubules in which stellate cells are expressing GFP (green) confirmed the co-localisation of dextran and GFP (yellow); DAPI, blue. Scale bars = 40 μm.

### Generality of the stellate cell model

The segregation of active cation transport to principal cells, and chloride and water flux to stellate cells, may confer selective advantages and could potentially extend to other insects. Stellate cells are more widely distributed than previously thought (15); and we have previously shown that fluorescently-labelled kinin (the neuropeptide that stimulates the chloride conductance (4, 18)) marks stellate-like cells in most advanced endopterygote insects (44), suggesting an ancient and conserved role. To probe the route of water flux in insects without the benefits of *Drosophila* transgenics, we applied the dextran flux labelling technique to a panel of insects selected to represent the major exo- and endopterygote Orders, so providing an initial view on the two cell model (Fig. 6). Among the exopterygote insects, dextran selectively labelled stellate-like cells of all insect Orders except the beetles, where an extensive network was observed. Significantly, kinin genes are almost never found in this Order (45), consistent with a lack of cell specialisation. In the more primitive exopterygotes, the story is more varied; although kinin had labelled the epithelium rather generally, this general pattern was seen in the locust, but not a cockroach. As a first approximation, therefore, the two-cell model, that links chloride flux, kinin stimulation and water flux, seems to have broad applicability across the higher insect; but further sampling of multiple species will be required.

**Fig. 6.**
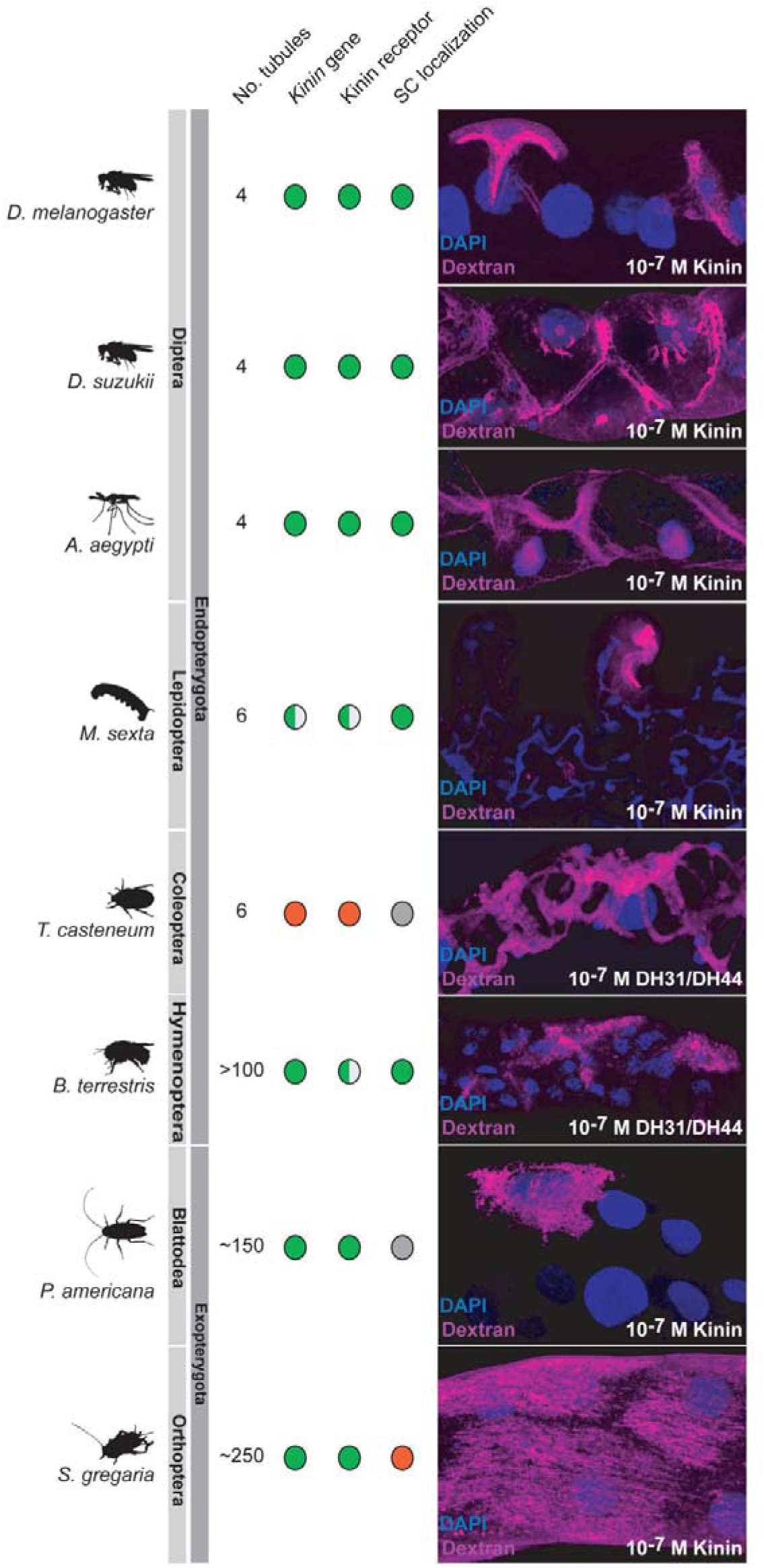
Dextran labelling is a tool to probe insect biodiversity. The dextran labelling protocol employed for *D. melanogaster* was applied to other insect species, selected from the major insect Orders of exopterygotes and endopterygotes. The distribution of known stellate cells and Kinin labelling (diagnostic of the route of chloride shunt conductance) is reproduced from (44).

### A revised model for a high-flux epithelium

The tubule shows a remarkable ability to secrete primary urine at very high rates; and together with other recent results, it is becoming clear that this success relies on the functional segregation of transport between different cell types (**Fig. 7**). The main, principal cell has long apical microvilli (40), each containing a mitochondrion (41), and loaded with proton-pumping V-ATPase and is thought to drive an exchanger from the NHA family to produce a net K^+^ flux. Basolaterally, the infoldings contain high levels of Na^+^, K^+^-ATPase (46), inward rectifier K^+^ channels (10, 11), and Na^+^/K^+^/Cl^−^ cotransporters (14). This metabolically active cell is likely the route for excretion of a wide range of solutes via ABC transporters and other organic solute transporters, many of which hare abundantly expressed in the tubule (24). The rarer stellate cells, by contrast have shorter microvilli and fewer mitochondria, but are the gatekeepers for the hormone-stimulated chloride shunt conductance (through basolateral ClC-a and apical secCl), and also for the passage of osmotically obliged water through basolateral Prip and apical Drip aquaporins. The metabolically active principal cell is thus sheltered from these very high, and potentially disruptive, fluxes of water.

**Fig. 7.**
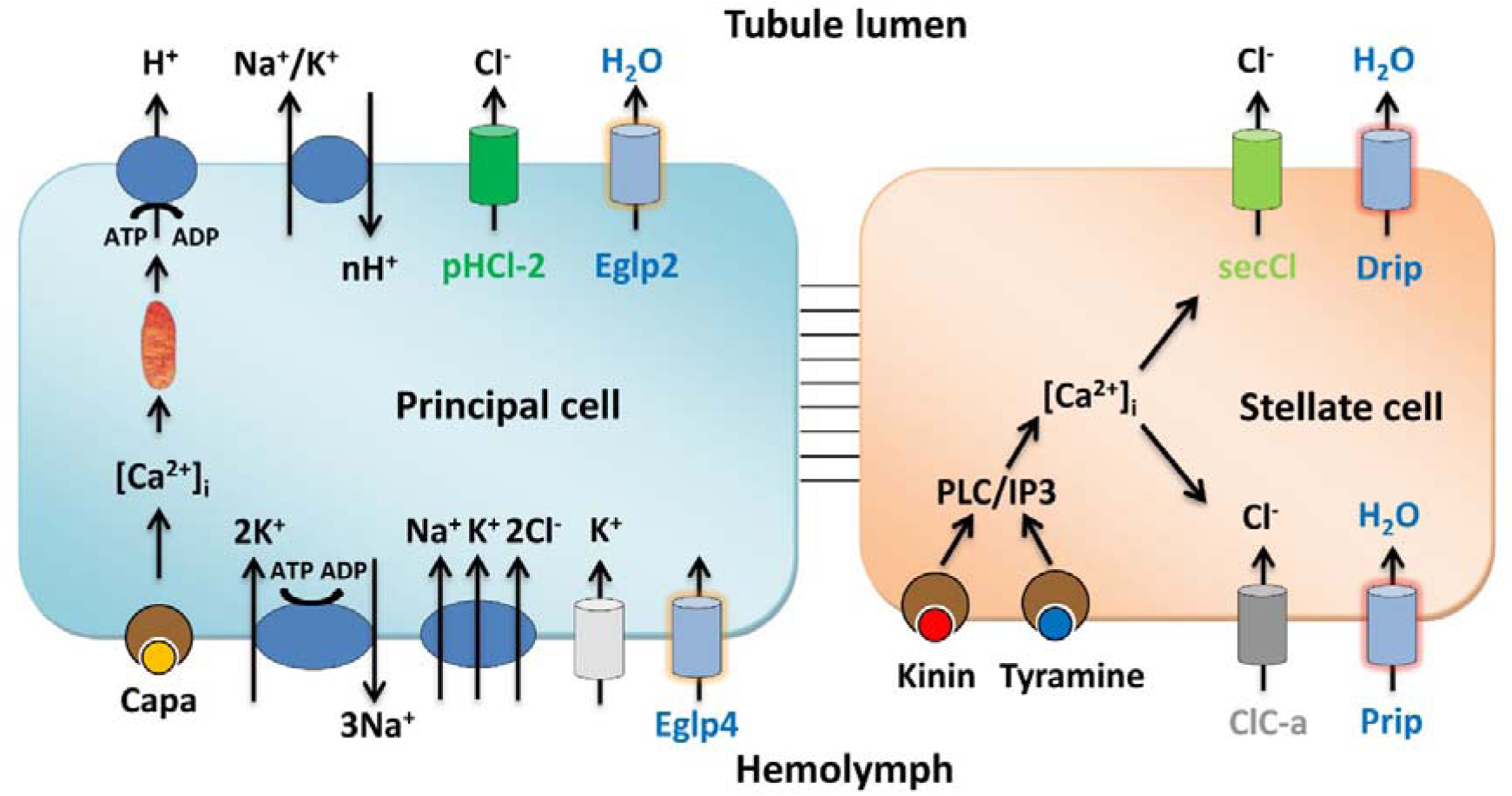
Model for tubule function. The mitochondria-rich principal cell is specialised for metabolically intensive cation and solute transport. The apical V-ATPase sets up a proton electrochemical gradient which drives net K^+^ secretion via NHA or NHE exchangers. Basolateral K^+^ entry is afforded by inward-rectifier K^+^ channels, Na^+^,K^+^ ATPase and an Na^+^,K^+^,2Cl^−^ cotransport. The resulting lumen positive potential drives a chloride shunt conductance, mainly via basolateral ClC-a and apical secCl channels in the stellate cell. The net transport of KCl drives osmotically obliged water, which is primarily via basolateral Prip and apical Drip in the stellate cells. In this way, the metabolically active principal cell is sheltered from the required high flux rates of water.

Given the severe consequences of unregulated fluid loss to a small terrestrial insect, it is not surprising that the tubule is under sophisticated neurohormonal control (47). Whereas cation pumping by the principal cells is under control of DH31 (48), DH44 (49) and Capa (50) neuropeptides, the stellate cells are independently controlled by the neuropeptide kinin (19, 51) and by the biogenic amine tyramine (17); both act indistinguishably through intracellular calcium (17, 52). The chloride shunt conductance is a known target of kinin and tyramine, as both rapidly collapse the lumen-positive potential (17, 18); however, it will be interesting to investigate whether one or both of this messengers have an independent action to regulate stellate cell aquaporins, perhaps through phosphorylation or recruitment to the plasma membrane.

This two-cell model is likely to be widely applicably through the higher insects, the endopterygotes, which include flies, butterflies and bees, in which a secondary cell type has been observed either directly (15), or by mapping an aquaporin (26), or by visualisation with fluorescently labelled kinin (the hormone which regulates chloride flux) (44), or by otherwise mapping the kinin receptor (53, 54). However, a universal model is unlikely, as most members of one higher insect Order (the Coleoptera) do not use kinin signalling (44); and in the lower exopterygote insects, such as crickets, there is no evidence for specialised secondary cells. The next challenge will be to map out the generality of this two-cell model, and its alternatives, across the tens of millions of species that makes up the insects.

## Materials & Methods

### Informatics

The MIP amino acid sequences were obtained from *Drosophila* gene database Flybase (flybase.org) and multiple sequence alignment was performed using Clustal Omega (www.ebi.ac.uk/Tools/msa/clustalo). Phosphorylation sites were analysed using GPS 3.0 algorithm (gps.biocuckoo).

### Drosophila Stocks and Rearing

Flies were reared at 22°C, 45% relative humidity on a 12:12 photoperiod on standard *Drosophila* media. The following lines (with original source) for this study were: Wild-type *D. melanogaster Canton-S* (Bloomington stock #1); *c724*-GAL4 (3) and ClC-*a-*GAL4 (VDRC #202625) driver lines specific to stellate cells, and used interchangeably in this study; *CapaR-*GAL4 driver line specific to principal cells (55, 56); *UAS-Drip*-*Venus* (21); dsRNA line directed against *Eglp2/CG17664* (VDRC #101847); *Eglp4/CG4019* (NiG-Fly stock #4019R-2); *Prip/CG7777* and *Drip/CG9023* respectively (NiG-Fly stock #7777R-2 and #9023R-2).

### Generation of Transformants

*UAS-Prip-Venus, UAS-Eglp2-Venus* and *UAS-Eglp4-Venus* were generated by PCR amplifying the coding sequence of the respective *Drosophila* MIP genes using DreamTaq green PCR master mix (Thermo Fisher Scientific) and the primer pairs listed in Table S1. ORF amplicons were cloned into pENTR donor vector (Invitrogen) and transferred to pTWV destination vector (DGRC) using Gateway® LR Clonase® II Enzyme mix according to manufacturing instructions (Thermo Fisher Scientific). Sequences integrity were confirmed by GATC Biotech and transgenic lines were generated by using standard methods for P-element mediated germ-line transformation (BestGene).

### Quantitative RT-PCR

For validation of tubule mRNA expression, qRT-PCR was performed using an ABI StepOnePlus Detection System (Applied Biosystems) with Brilliant III Ultra-Fast SYBR Green QPCR master mix (Agilent, UK) and the primer pairs listed in **Table S1**. Data were normalized against the *rpl32* standard and expressed as fold change compared to controls ± SEM (n = 3).

### Antibody Production and Immunohistochemistry

Antigenic peptides were identified using Abdesigner software (57). Rabbit anti-peptide antibodies were raised against the Drip epitope (CFKVRKGDDETDSYDF), Prip epitope (CNEASEKYRTHADERE), Eglp2 epitope (CSEVDETTMSTKRTSE), and Eglp4 epitope (CTSNEKLRQLEDVQLS) by Genosphere Biotechnologies (Paris, France).

Malpighian tubules from 7-day old flies were dissected in Schneider’s medium (Thermo Fisher Scientific) and transfer to poly-L-lysine (Sigma-Aldrich)-covered 35-mm glass-bottomed dishes (MatTek Corporation) in PBS, fixed in 4% (w/v) paraformaldehyde in PBS for 30 min at room temperature, washed in PBT (PBS + 0.05% (v/v) Triton X-100), and then blocked in 10% (v/v) normal goat serum (Sigma-Aldrich) in PBT. IgG purified rabbit anti-Drip/Prip/Eglp2/Eglp4 peptides (dilution 1:1000) were used. Alexa Fluor 488/564-conjugated affinity-purified goat anti-rabbit antibodies (Thermo Fisher Scientific) were used at a concentration of 1:1000 for visualisation of the primary antiserum. Incubations in the primary and secondary antibodies were performed overnight at 4°C. Tubules were incubated with the nuclear stain DAPI (1 μg mL^−1^; Sigma-Aldrich) for 1 min and in some cases Rhodamine/Alexa-633 coupled phalloidin (1:100; Thermo Fisher Scientific) in PBT for a minimum of 30 min. Samples were washed repeatedly in PBS before being mounted in Vectashield (Vector Laboratories Inc). Confocal images were taken using an LSM 880 inverted microscope (Zeiss) and processed with Zen black/blue software (Zeiss, Oberkochen, Germany) and Adobe Photoshop/Illustrator CS 5.1.

### Western Blotting

For each fly line, Malpighian tubules from >50 flies were dissected under Schneider’s *Drosophila* medium (Thermo Fisher Scientific) and were transferred to 100 μl RIPA buffer (150⍰ mM NaCl, 10⍰ mM Tris-HCl pH 7.5, 1⍰ mM EDTA, 1% Triton X-100, 0.1% (wt/v) SDS) with 1 μl of protease inhibitor cocktail (Sigma-Aldrich). Samples were homogenized using a Microson XL2000 sonicator (Misonix Inc. NY, USA), and centrifuged (13,000 rpm) at 4°C for 10⍰ min. Protein concentrations were measured using the Bradford Protein Assay (Bio-Rad Technologies). Approximately 20⍰ μg protein from each sample was electrophoresed on a NuPage 4–12% Bis-Tris gel and blotted onto nitrocellulose membrane using the Novex system (Thermo Fisher Scientific). Blots were stained with Ponceau S and probed with IgG purified rabbit anti-Drip/Prip/Eglp2/Eglp4 antibodies (1 μg mL^−1^) and developed by electrochemiluminescence assay using ECL^™^ horseradish peroxidase linked anti-rabbit IgG (1:2000; Amersham Biosciences).

### Fluid Secretion

Secretion assays were performed as described previously (2). Malpighian tubules from 7-day-old adult female flies were dissected under Schneider’s insect medium (Thermo Fisher Scientific) and isolated into 10 μl drops of a 1:1 mixture of Schneider’s medium : *Drosophila* saline. Intact tubules were left to secrete for approximately 30 min before starting the experiment. Secretion rates were measured every 10 min; after 30 min of baseline readings, the diuretic peptide *Drosophila* kinin (DK) and capa-1 were added to 10^−7^M, and secretion rates measured for a further 30 min. Data are plotted as mean ± SEM (n > 7).

### Dextran Labelling

Individuals were lightly anaesthetised using either C0_2_ or ice and their Malpighian tubules dissected in Schneider’s *Drosophila* medium (Thermo Fisher Scientific). Dissected tissues were then pre-incubated for 10-20 min at room temperature in a solution of 1:1 Schneider’s:PBS with neuropeptides (e.g. Kinin, DH31, DH44) present at a concentration of 10^−7^ M (stimulated) or with no neuropeptides (non-stimulated). The dissected tissues were then transferred to fresh Schneider’s:PBS solution containing 0.2% dextrans (40 kDa or 70 kDa; Thermo Fisher Scientific) conjugated to a specified fluor, for 2 – 5 min at room temperature. Tissues were fixed for 10 min in 2% (w/v) paraformaldehyde, stained with 1 µg/ml DAPI (Sigma) for 2 min, transferred to poly-L-lysine (Sigma)-covered 35-mm glass-bottomed dishes (MatTek Corporation) in PBS, and imaged using a Zeiss LSM 880 confocal microscope (Zeiss).

### *Xenopus* Oocyte Assays

cDNAs for *Drosophila* AQPs (Drip, Prip, Eglp2, Eglp4), human AQP4 and mefugu AQP8 were cloned into pGEMHE, a plasmid optimized for cRNA expression in *Xenopus laevis* oocytes. cRNA synthesis, oocyte injections (10 ng/oocyte) and oocyte care were performed as previously (58). To ensure basic water channel-activity before detailed analysis, oocytes expressing AQPs were placed in distilled water and time for swelling and ultimately bursting noted.

To calculate permeability to water (osmotic), glycerol, mannitol and urea, we used a Zeiss Lumar, ZEN 2.0 and a four well-perfusion chamber (1-1.5 ml) and acquired images every 5 s for 10 min. For osmotic water permeability, we diluted ND96 (200 mOsm) to 70 mOsm. Permeability for glycerol, mannitol and urea, were assessed by exposing oocytes to 200 mM solute (2 mM HEPES, pH 7.5) from ND96. Solutions were perfused into the chamber (full solution change in 20 s) beginning at 45-50 seconds into the experiment. To prevent contamination effects, i.e., not returning to baseline volume, oocytes were only exposed to one osmotic or solute challenge. The experiments were repeated for each of the different substrates, and oocytes from at least three different donor *Xenopus* were used.

Water permeability (P_f_) was calculated as before (59, 60):

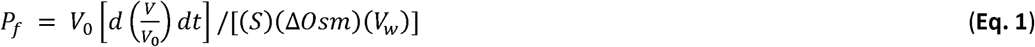

#### Data analysis

Experiments were analyzed using a custom macro in FIJI. The image files were first converted to an 8-bit image then made binary. The analyze particles function was used to measure the major (a) and minor (b) diameter of all four oocytes (3µm per pixel). The 3^rd^ axis (c) for ellipsoid volume was calculated as the average of the major and minor axes. Ellipsoid surface area is calculated as:

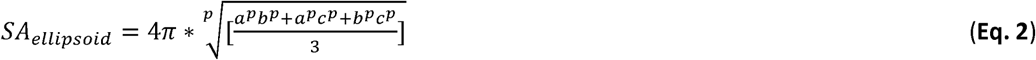

where p ≈ 1.6075. This SA_ellipsoid_ is then multiplied by 8 to account for the SA convolutions of the oocyte. Finally, Permeability of solute (P_solute_) is calculated:

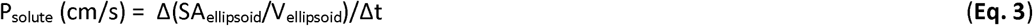

where Δt= each 20 s (4 point) interval after solute addition. P_solute_ (cm/s) was maximal by 120s after change to 200 mM solute. We then used a 4 point rolling average 120 s after the test solute as the reported value P_solute_ (cm/s) for each oocyte. Number of injected oocytes used for each experiment are presented in **Table S2.**

### Desiccation Stress Assay

7-day old male and female flies of specified genotype were anaesthetised briefly with CO_2_ and placed in groups of 30 in empty vials (no food or water), and the open end of the tube was sealed with parafilm (Bemis, NA, USA). Flies were counted until 100% mortality was reached and data expressed as % survival ± SEM (n = 3).

### Statistical Analysis

GraphPad Prism 7.0 software (GraphPad Software Inc., CA, USA) was used for statistical analysis and generation of graphs. For fluid secretion analysis, a two-tailed Student’s t-test, taking *P* = 0.05 as the critical value (for two independent groups: basal versus stimulated), was used. For mRNA level quantification, one-way analysis of variance (ANOVA) followed by Tukey’s multiple comparisons of means with a significance level of *P* < 0.05 (for three independent groups) was used. For measurement of oocyte water and solutes permeability, we used an ANOVA analysis for each group with a significance level of *P* < 0.05. For survival curves obtained in desiccation assays, significance was assessed by the log-rank (Mantel-Cox) test. Log-rank tests were conducted for each pairwise comparison.

## Acknowledgments

We thank Vilija Lomeikaite, Keith Graham and Leonardo Beltránfor experimental assistance. Fly lines were obtained from the VDRC RNAi stocks (Vienna Drosophila Resource Center, Austria), Bloomington Stock Center (Indiana University, Bloomington, USA) and Fly Stocks of National Institute of Genetics (Japan). This work was funded by the Biotechnology and Biological Sciences Research Council (UK) grants BB/L002647/1 (SD/JATD/ST) and BB/P008097/1 (SD/JATD/ST) and National Institutes of Health Grants DK100227, DK101405 (MFR).

