## Supplementary data for "When two cells are better than one: specialized stellate cells provide a privileged route for uniquely rapid water flux in *Drosophila* renal tubule"

### Supplementary Figures

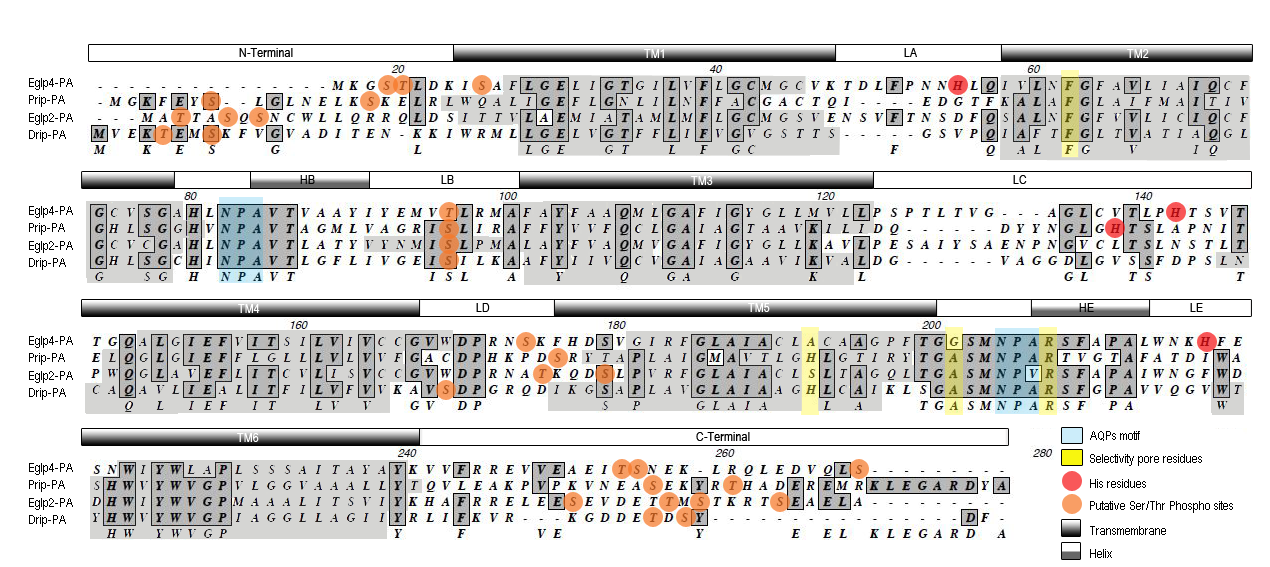

**Fig. S1. Clustal Omega alignment of the amino acid sequences of *Drosophila melanogaster* aquaporins (Drip and Prip) and aquaglyceroporins (Eglp2 and Eglp4).** MIP proteins consist of two tandem repeats, each of which has three membrane spanning *α*-helices and a pore forming loop with a signature Asp-Pro-Ala (NPA) motif (in blue). Highlighted in orange are the putative phosphorylation sites; in red are the Histidine residues involved in pH and divalent cations sensitivity; in yellow are the residues involved in pore selectivity.

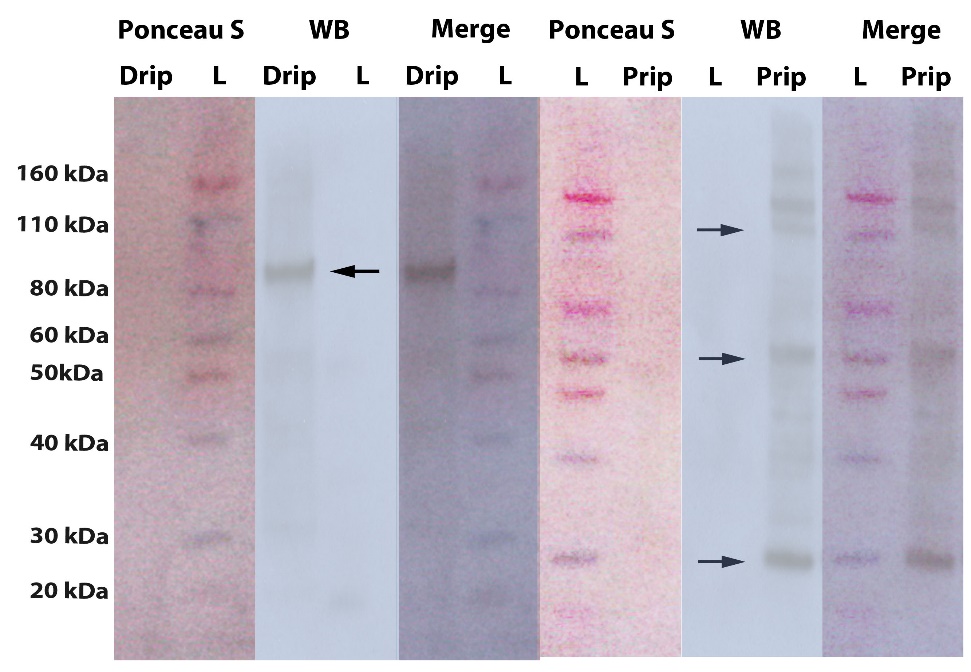

**Fig. S2. Western blot of Malpighian tubules protein homogenates probed with Drip and Prip antibodies.** Ponceau S staining and western blot of tubule protein samples of Drip and Prip antibodies. The predicted molecular mass of Drip and Prip monomer is 25.5 and 29 kDa respectively and specific bands corresponding to either monomer, and higher molecular weight bands corresponding to dimer and/or tetramer were detected (arrows).

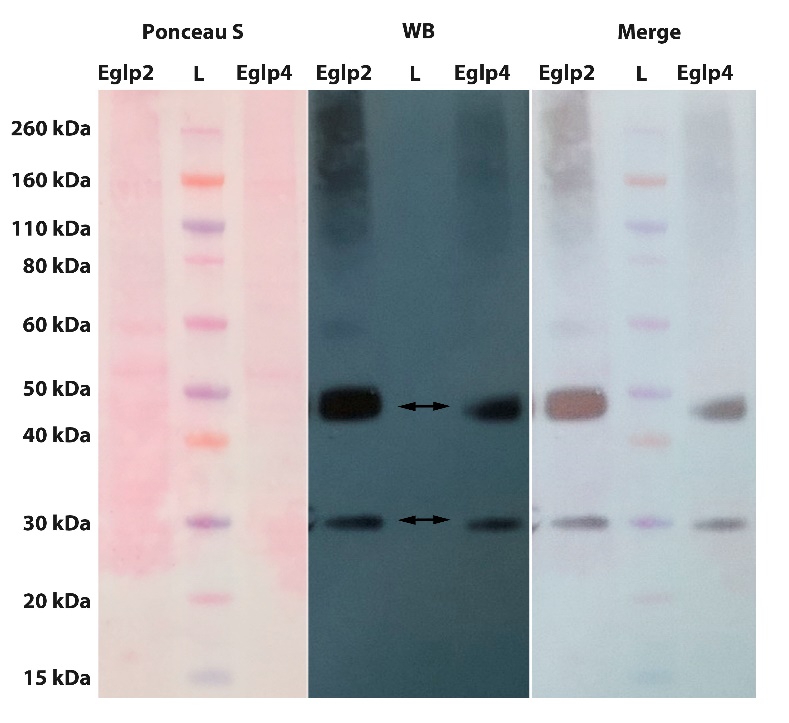

**Fig. S3. Western blot of Malpighian tubules protein homogenates probed with Eglp2 and Eglp4 antibodies.** Ponceau S staining and western blot of tubule protein samples of Eglp2 and Eglp4 antibodies. The predicted molecular mass of Eglp2 and Eglp4 monomer is 28.8 and 30.9 kDa respectively and specific bands corresponding to either monomer and dimer were detected (double arrows).

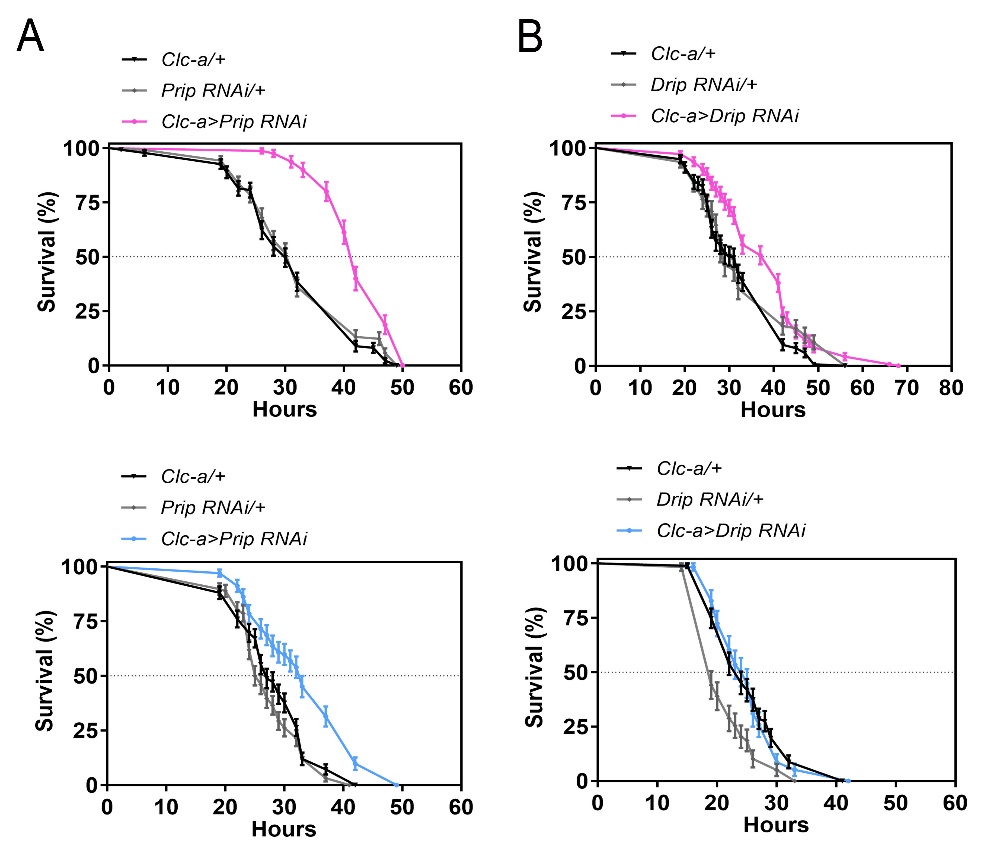

**Fig. S4. Compromising the stellate route for water flux increases survival on desiccation**. Female (top) or male (bottom) flies were subjected to desiccation stress. (*A*) Reduced Prip levels in stellate cells (*ClC-a*-GAL4>*UAS-Prip RNAi*) alter survival of desiccated flies. Desiccation resistance was significantly higher after knockdown of *Prip* in stellate cells compared to controls (P < 0.001 against both controls; Log rank test, Mantel-Cox). (*B*) Reduced Drip levels in stellate cells have a similar effect to Prip downregulation in female flies under desiccation conditions.

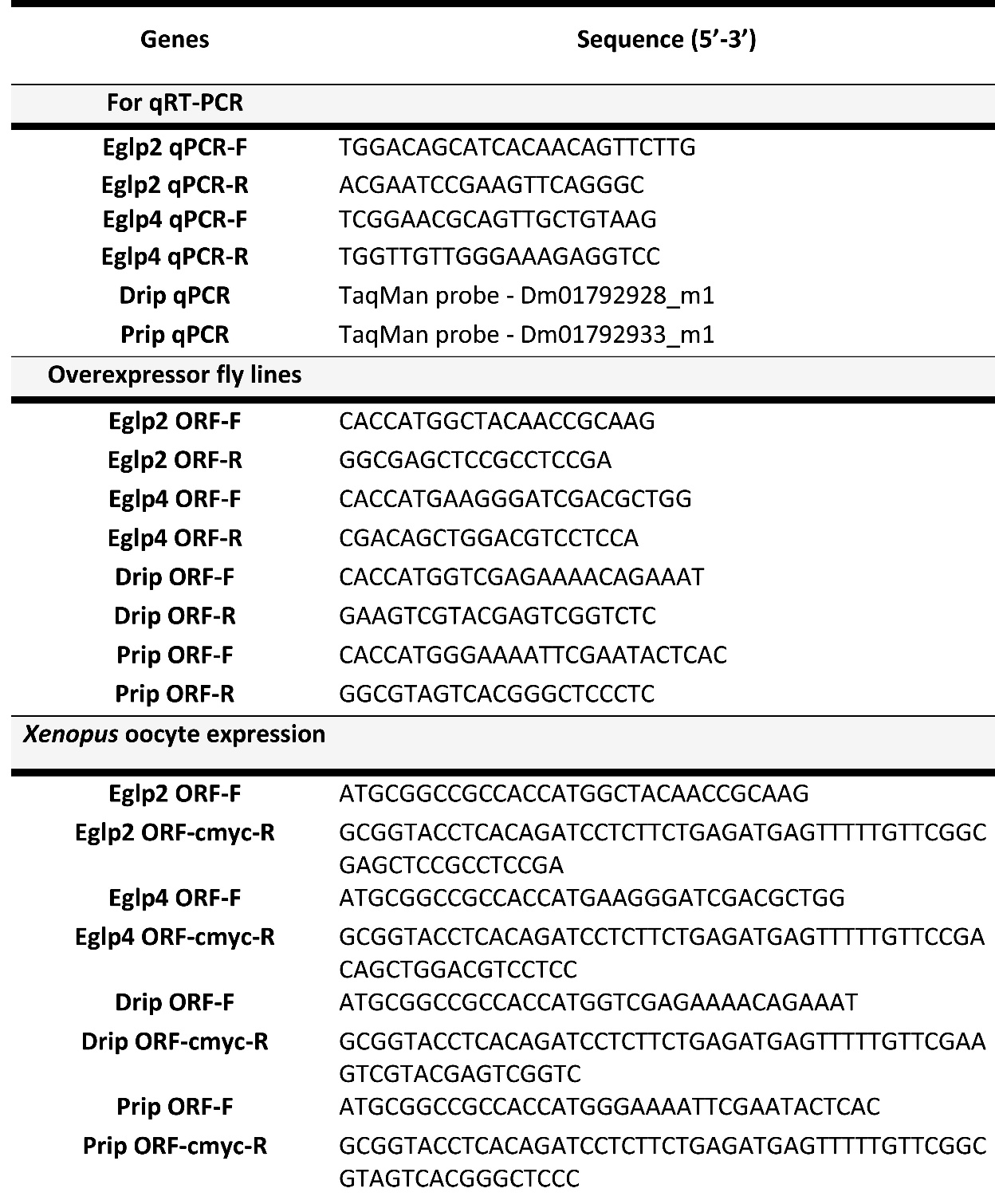

**Table S1. Primer sequences used in this study.**

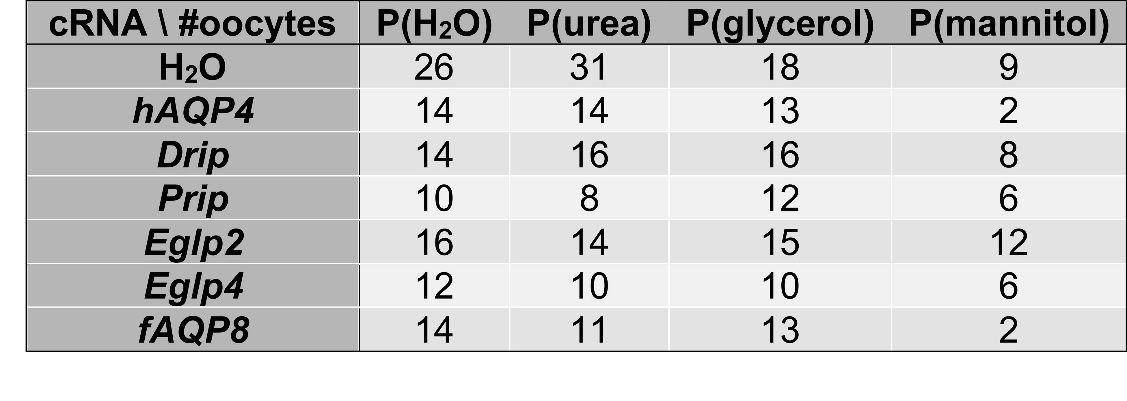

**Table S2. The table indicates the total number of oocytes for each MIP permeability experiments.**
